# On X-ray Sensitivity in Xenopus Embryogenesis

**DOI:** 10.1101/2025.03.25.645254

**Authors:** Hugo M. Santos, Jose G. Abreu, Leonid Peshkin

## Abstract

We examined the effects of X-ray irradiation on Xenopus laevis, focusing on pre- and post-fertilization exposure. We applied X-ray doses of 10, 50, 100, 250, and 500 Gy. Fifty percent of the 360 eggs irradiated at 250 Gy failed to fertilize, while fertilized eggs developed normally until the gastrula stage. Doses ranging from 10 to 250 Gy caused developmental anomalies. High mortality rates were observed at doses of 100 to 500 Gy. Post-fertilization irradiation at 50 to 100 Gy resulted in 100% lethality, while exposure to 10 Gy led to only 13% lethality, although both exposure levels produced similar types of developmental anomalies compared to pre-fertilization irradiation. This study highlights how the timing and intensity of exposure critically affect embryo viability, especially during the sensitive stages of fertilization and gastrulation. We establish the necessary and sufficient dosage to further investigate the molecular mechanisms of X-ray damage to DNA and protein.

**Highlight:** High doses of electromagnetic radiation pre- and post-fertilization produce different developmental outcomes in Xenopus laevis.

## Description

X-rays primarily interact with human and animal cells through the ionizing radiation they emit, which can lead to various effects depending on the dose and duration of exposure. High ionization levels can remove tightly bound electrons from atoms, creating ions. This process can result in DNA breakage, impair the cellular repair mechanisms, and ultimately cause cell death (Saini and Gurung, 2025; Huang and Zhou, 2021). In addition to DNA damage, X-rays can directly or indirectly affect proteins, which are critical for cellular structure and function.

Ionizing radiation induces protein damage through direct ionization and ROS-mediated oxidation (e.g., hydroxyl radicals), leading to carbonylation, fragmentation, and cross-linking of amino acid side chains (Hall & Giaccia, 2018; Dalle-Donne et al., 2006). This disrupts enzymatic activity, signal transduction pathways, and structural integrity, exacerbating cellular dysfunction. Accumulated damage overwhelms proteostasis systems (e.g., ubiquitin-proteasome, chaperones), triggering ER stress, misfolded aggregates, and apoptosis (Malhotra & Kaufman, 2007). Carbonylated proteins serve as oxidative stress biomarkers linked to neurodegeneration, aging, and radiation-associated pathologies (Stadtman & Levine, 2003; Azzam et al., 2012). Gametes and zygotes are particularly radiosensitive compared to somatic cells due to radiation-induced DNA damage, interference with the cell cycle, and differentiation during these phases. This presents a convenient model to induce and study protein damage and aggregation in a controlled way.

Amphibians, including Xenopus laevis, have been utilized as models to study the effects of a wide range of radiation doses during embryonic development, pre-metamorphosis, and adult stages (Hamilton, 1967; Ijiri, 1979; Landreth and Dunaway, 1974; Anderson et al., 1997; Dong et al., 2010; Stark et al., 2015; Carotenuto et al., 2016). These studies generally indicated lethal doses at various stages of the amphibian life cycle. Research by Hamilton in the late 1960s demonstrated that haploid Xenopus embryos are more sensitive to X-ray exposure than their diploid counterparts (Hamilton, 1967). Hamilton attributed this difference in X-ray dose response to genetic factors related to chromosome number. However, his study raises the possibility that DNA damage alone may not fully account for the observed results, suggesting that protein damage could also play a significant role.

While these studies have highlighted how amphibians respond to radiation, further investigation is needed to understand their resistance to extreme doses. We decided to investigate the effects of pre- and post-fertilization radiation on the survival and embryonic development of Xenopus laevis. We applied various X-ray doses: 10, 100, 250, and 500 Gy, to the Xenopus eggs, and then fertilized them 1h after irradiation to observe the effects. We applied 10, 50, and 100 Gy to newly fertilized embryos (Fig. 1A).

**Figure 1.**
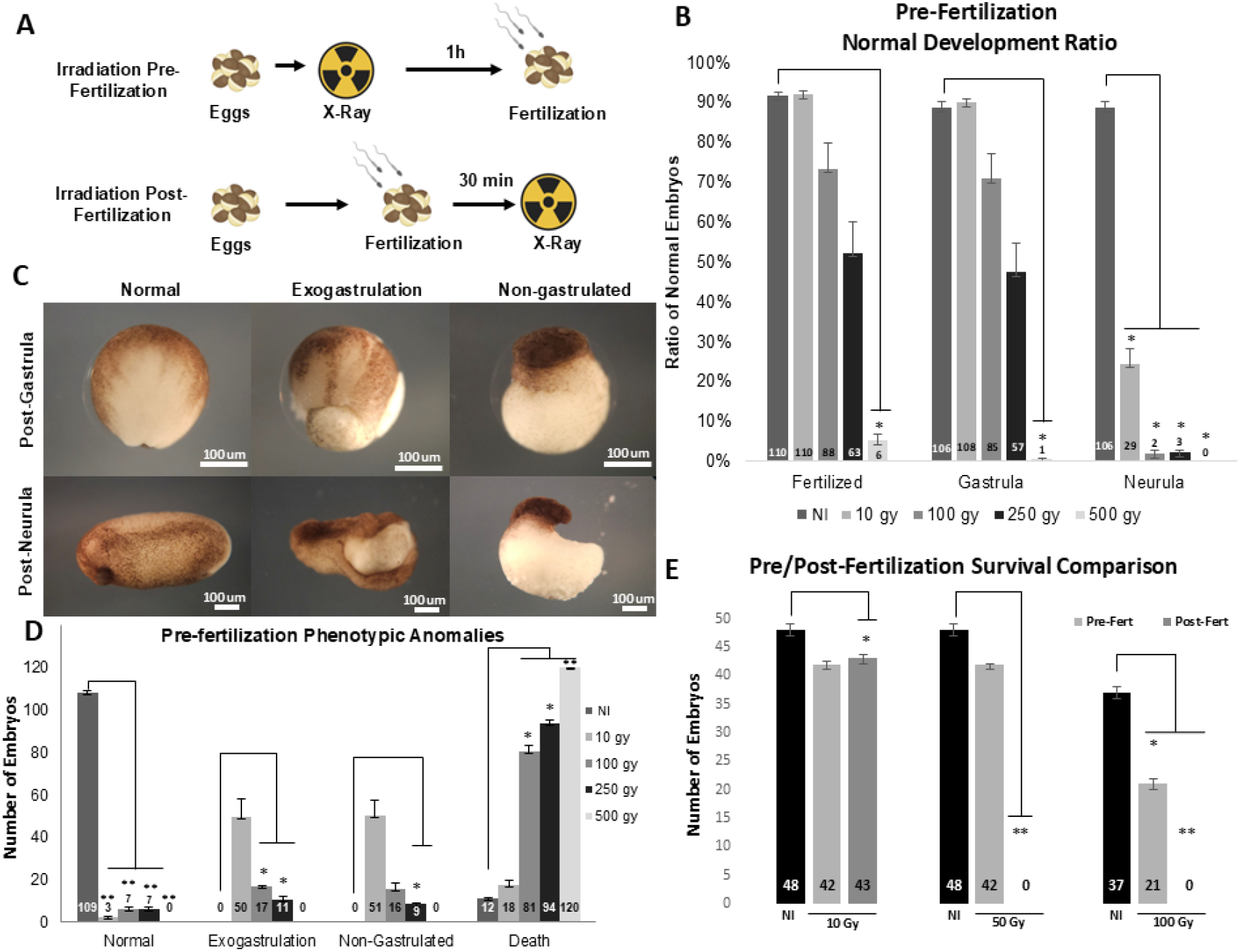
X-Ray Impact on Xenopus laevis development pre- and post-fertilization. (A) Two alternative experimental designs: irradiation pre- and post-fertilization. (B) Bar plot of the average normal development of eggs irradiated with 10, 100, 250, and 500 Gy pre-fertilization. P values: t-test, n=3, with 120 eggs per condition in triplicates from 3 different females (total 360 per condition). Conditions significantly decreased compared to control (*: two-tailed paired t-test P < 0.05). (C) Sample embryos illustrate the phenotype classes observed during X-ray exposure: normal, exogastrulation, non-gastrulation, and death. (D) Distribution of phenotypes observed in C for different pre-fertilization X-ray exposures. P values: t-test, n=3, with 120 eggs per condition in triplicates from 3 different females (total 360 per condition). Conditions significantly decreased compared to control (*: two-tailed paired t-test P < 0.05; **: t-test P < 0.001). Subfigures B and D show quantification from two independent experiments. (E) Average survival of late neurula embryos from pre- and post-fertilized irradiated groups at 10, 50, and 100 Gy. Each group started with 50 initial eggs or zygotes in triplicates per condition of two independent females (300 embryos per condition tested). P values: t-test, n=2 of 150 embryos per condition. Conditions significantly decreased compared to control (*: two-tailed paired t-test P < 0.05; **: t-test P < 0.001).

During the initial evaluation, we observed two key effects of radiation: a significant reduction in fertilization rates, particularly at the highest doses of 250 and 500 Gy, where fertilization rates dropped to approximately 50% and 5% on average, respectively (Fig. 1B, first column table). We then noticed that the fertilized eggs showed normal development up to the blastula stage. We also observed an increase in the average number of embryos that fail to gastrulate as the radiation dose increases (Fig. 1B, second table column). From gastrula to neurula, the radiation effect was dramatic; at just 10 Gy, an average of 76% failed to reach the neurula stage, and at 500 Gy, no embryo succeeded in transitioning from gastrula to neurula (Fig. 1B, third table column). In another independent experiment, we evaluated four phenotypic categories observed in pre-fertilized radiated and non-radiated eggs (Fig 1C and D). In non-radiated conditions, an average of 91% of embryos presented normal development (Fig 1C and D). Conversely, several anomalies were observed at radiation doses from 10 to 500 Gy, including death, exogastrulation, and non-gastrulation (Fig 1C and D). High mortality was observed mainly in the groups that received 100, 250, and 500 Gy, presenting a mortality rate of 67.5%, 79%, and 100% embryo on average, respectively (Fig. 1D). Only an average of 2.5-6% of the irradiated embryos managed normal development.

For the post-fertilization experiment, we based our design on Hamilton’s study, which applied a maximum dose of 50 Gy to diploid embryos (Hamilton, 1967). Consequently, we subjected 50 zygotes per condition, in triplicate from two distinct females to radiation doses of 0, 10, 50, and 100 Gy in order to compare the average survival of radiation exposure pre-fertilization to post-fertilization. We counted the number of surviving embryos at the late neurula stage (Fig 1E). No embryos reached the neurula stage at the 50 and 100 Gy doses (Fig 1E, dark grey bars). At the 10 Gy dose, the average number of surviving neurula embryos was roughly similar but significantly lower than that of the non-irradiated embryos (Fig 1E, first three bars).

This study builds on and further elucidates X-rays’ effects, expanding on Hamilton’s foundational work (1967). By revisiting this topic, we introduced a pre-fertilization irradiation condition and observed that egg infertility increased proportionally with higher radiation doses. In contrast, eggs exposed to lower radiation levels exhibited a higher rate of phenotypic abnormalities but lower lethality during embryonic development. Our findings align with Hamilton’s (1967) observations, showing that pre- and post-fertilization irradiated embryos can survive until the blastula stage without anomalies. However, as gastrulation begins, significant anomalies emerge, often leading to embryonic death. Notably, pre-fertilization irradiation demonstrated greater damage tolerance than post-fertilization conditions. This finding strongly suggests that the new sperm content (proteins and genetic material) counteracts to some degree the egg damage caused by X-ray irradiation from 10 to 250 Gy, which is an extreme dose.

In summary, this study developed a reliable methodology for challenging chromosomes and cellular constituents, such as proteins, through X-ray radiation in Xenopus eggs and embryos.

## Methods

### Animal Preparation

To prepare female animals for ovulation, they were primed with 100 IU of Human Chorionic Gonadotropin (hCG) (Chorulon 5000 IU – Merck A002377) 72 hours before fertilization. The night preceding fertilization, each animal received a booster injection of 750 IU of hCG and was kept in individual containers at 18°C to initiate ovulation.

On the day of fertilization, male testes were surgically extracted according to protocol. Green, Sherril L. The Laboratory Xenopus. 1^st^ Edition. 2010. The extracted testes were then preserved in a 1X MMR (NaCl 1M, KCl 20 mM, MgSO4 10 mM, CaCl2 Dihydrate 20 mM, HEPES 50 mM) solution at 4°C.

### Egg Collection

Approximately 12 hours following the hCG boost, the females were transferred to containers containing a 1X High Salt Barth solution (NaCl 1.095 M, KCl 0.010 M, NaHCO3 0.024 M, MgSO4-7H2O 0.008 M, HEPES 0.100 M, Ca(NO3)2 – 4H2O 0.003 M, CaCl2-2H2O 0.004 M). After about 90 minutes, eggs were collected for either fertilization or irradiation.

### Fertilization Process

After draining the High Salt Barth solution, eggs were fertilized using macerated testes in 1X MMR. The sperm-egg mixture was homogenized for 4 minutes, followed by adding 0.1X MMR. Cortical rotation occurred within 20 minutes. A 2% cysteine solution (Millipore 1.028838.0100) was added, and embryos were agitated 5 times every 2 minutes, rinsed with 0.1X MMR, and cultured at 18°C.

### Irradiation Procedure

Irradiation was carried out using a Rad Source RS 1800 Q X-ray machine. A mandatory 3-hour cooling period between sessions is required in experiments with 500 Gy. Each irradiation session was limited to no more than 20 minutes. Doses of 100, 250, and 500 Gy were administered on shelf 2 of the instrument (9.5 cm from the source), with samples centralized to ensure optimum radiation exposure. A 10 Gy irradiation was conducted on the instrument’s floor (21.5 cm from the source), adhering to the same distribution principles. All samples were irradiated in 35 mm Petri dishes (Falcon 353001) containing 50 to 120 eggs or embryos each. Eggs irradiated before fertilization were suspended in a small amount of 1X Barth High Salt solution for preservation. Embryos were irradiated in a 0.1X MMR solution immediately after the cysteine washing step and before the first cleavage stage.

## Reagents

| Reagent | N° catalog | Available From |
| --- | --- | --- |
| Chorulon 5000 IU (hCG) | A002377 | Merck |
| cysteine | 1.028838.0100 | Millipore |
| Falcon 35 mm Petri dishes | 353001 | Corning |
| NaCl | SX0420-5 | Millipore |
| KCl | 7300-500GM | Millipore |
| MgSO<sub>4</sub>·7H<sub>2</sub>O | 230391-500G | Sigma |
| HEPES | H-400-1 | Goldbio |
| Ca(NO<sub>3</sub>)<sub>2</sub>·4H<sub>2</sub>O | A16645.36 | Thermoscientific |
| CACL<sub>2</sub> | C8106-1KG | Sigma |
| CACL<sub>2</sub>·2H<sub>2</sub>O | C3881-500G | Sigma |

